# Social selection is density dependent but makes little contribution to total selection in New Zealand giraffe weevils

**DOI:** 10.1101/2021.02.06.430048

**Authors:** David N Fisher, Rebecca J LeGrice, Christina J Painting

## Abstract

Social selection occurs when traits of interaction partners influence an individual’s fitness and can fundamentally alter total selection strength. Unlike for direct selection, however, we have little idea of what factors influence the strength of social selection. Further, social selection only contributes to overall selection when there is phenotypic assortment, but simultaneous estimates of social selection and phenotypic assortment are rare. Here we estimated social selection on body size in a wild population of New Zealand giraffe weevils (*Lasiorhynchus barbicornis*). We did this in a range of contexts and measured phenotypic assortment for both sexes. Social selection was mostly absent and not affected by sex ratio or the body size of the focal individual. However, at high densities selection was negative for both sexes, consistent with competitive interactions based on size for access to mates. Phenotypic assortment was also density dependent, flipping from positive at low densities to negative at high densities. However, it was always close to zero, indicating negative social selection at high densities will not greatly impede the evolution of larger body sizes. Despite its predicted importance, social selection may only influence evolutionary change in specific contexts, leaving direct selection as the dominant driver of evolutionary change.

## Introduction

Selection is an important concept in evolutionary biology, describing the link between traits and fitness. Typically, selection is characterised as a selection gradient or covariance between the trait of a focal individual (e.g. its body size) and a measure of fitness (e.g. the number of adult offspring it has over lifetime; [1]). This “direct” selection helps us understand the ultimate functional value of traits and predict how they might evolve. Further, direct selection is known to vary across space [2], time [3], and with ecological conditions [4,5], helping to generate the biodiversity of the natural world. Other forms of selection are possible, however. For instance, when organisms interact with others, such as by competing for access to resources or cooperating to raise young, they can influence each other’s fitness. The link between a partner’s traits or the traits of group mates and a focal individual’s fitness is known as “social” selection [6]. Social selection may not align with direct selection (see Table 3 of [7]), which can alter the direction of trait evolution [8]. For instance, “selfish” traits may increase the fitness of an individual that bears them but be costly when displayed by group mates. Conversely, “altruistic” traits may be costly for the individual that displays them but be beneficial when possessed by group mates. Social selection can therefore be expected to alter evolutionary change and trait optima away from that expected solely under direction selection, making it a fundamentally important evolutionary parameter [6,9,10].

Social selection alone cannot alter evolution, however. For social selection to contribute to total selection, and therefore evolutionary change, there must be non-zero phenotypic assortment among interacting individuals [6]. Phenotypic assortment describes the covariation between the traits of an individual and the traits of those it interacts with. Positive assortment indicates that individuals with similar traits interact e.g., aggressive individuals interact with other aggressive individuals. Negative assortment on the other hand indicates that individuals with dissimilar traits interact e.g., resource producing individuals interact with resource consuming individuals. Assortment has been often documented in groups of animals, and typically found to be positive (in male great tits, *Parus major*, [11], Chacma baboons, *Papio ursinus*, [12]; guppies, *Poecilia reticulata*, [13]). However, not all measures of assortment are equal, and only the interactant covariance (the covariance between an individual’s traits and the mean trait value of those it interacts with) is correct for use in models of total selection [14]. Unfortunately, estimates of this parameter in natural populations are rare, especially alongside estimates of social selection (but see: [15]). Therefore, despite its predicted importance, we have very little knowledge of how social selection contributes to total selection in natural populations.

Alongside limited knowledge of social selection’s contribution to overall selection, we also have little data on the contexts where social selection is strongest (but see: [16,17]). Direct selection is known to vary based on demographic parameters such as population density [18] and sex ratio [19], as well as depending on other individual characteristics (correlational selection; [20]). A lack of knowledge of the conditions where social selection is strongest completely hampers our ability to predict how it may shape different populations differently, and therefore generate diversity. We aimed to fill gaps in our knowledge surrounding the contribution of social selection to total selection, and the conditions it is strongest, in a wild population of New Zealand giraffe weevils (*Lasiorhynchus barbicornis*; Coleoptera: Brentidae). Both sexes are extremely variable in size [21–23], males bear an elongated rostrum used as a weapon during contests for mates [24], and body length is under positive linear, but not quadratic, selection in males and females [25]. As giraffe weevils form aggregations on trees and compete for access to mates, we predicted that social selection for body size would be negative, where the presence of larger rivals reduces a focal individual’s fitness (following [26]). Further, we predicted that this social selection would be more negative at high densities and when the individual was of the more common sex (i.e., a male in a male-biased population), as these are conditions when they might be competing most fiercely for access to mates. We also predicted that smaller males would be less affected by the body size of rivals, as they can readily switch between fighting with similar-sized rivals and “sneaker” tactics that allow them to gain copulations without direct competition [27]. Finally, following previous work which found positive size-assortment among mating pairs [25], we predicted that there would be positive assortment for body size in the individuals present on trees in both sexes, causing social selection to reduce the overall strength of selection on body size.

## Methods

### Data collection

The giraffe weevil population we studied resides in Matuku Reserve (36° 51.92⍰S, 174° 28.32⍰E), an area of native coastal broadleaf forest west of Auckland, New Zealand. We located aggregations of adult giraffe weevils on karaka trees (*Corynocarpus laevigatus*), which were subsequently used for behavioural observations. The observations and data collection used in the current study are described in full in a previous study [25] with the data available online [28], but we briefly outline them again here.

To determine variation in mating success among males and females of different sizes we conducted daily observations for one hour at three different trees that housed giraffe weevil aggregations. Observations took place over two periods between November 22 and December 22, 2013 (31 days, N = 120 females, 132 males), and January 22 to February 23, 2014 (33 days, N = 301 females, 366 males). For the analysis we excluded individuals only seen once, and those who were first seen in the last week of each observation period (following [25]). This left a dataset of 1234 records of 155 different females and 236 different males. At least two hours prior to observations each day, we removed all giraffe weevils from the tree for measurement and marking. We measured total body length (tip of mandibles to distal end of elytra) using digital callipers to nearest 0.01 mm. We also measured weapon size and other morphological traits, but these are all very highly correlated with body length, while body length includes the rostrum (the weapon) and is likely under fecundity selection in females, hence is an appropriate trait to use for our analysis [22]. We then painted individuals on the pronotum and elytra with a unique colour combination using five Queen bee marking paints (Lega, Italy) for identification before being released to the point of capture on the tree [29]. We observed all giraffe weevils present on each of the three study trees for one hour on each day of the observation period between 0800 h and 1800 h. We stood at least one metre from the tree and used close range binoculars (Pentax, Papilio) to avoid disturbing the weevils. During each observation, we recorded the identification of all giraffe weevils present on the tree that day as well as all matings. After observations, we thoroughly searched the tree to check for any individuals that had been inactive or hiding in cracks or under leaves, and we gave these individuals a mating frequency of zero. We conducted no observations on days of heavy rainfall because giraffe weevils are inactive during this time, resulting in two non-consecutive days being missed in the first observation period, and three non-consecutive days during the second.

### Data analysis

To assess the strength of social selection, we fitted a series of generalised linear mixed-effect models using the R package “glmmTMB” [30]. For all models we mean-centred each continuous predictor variable and divided by its standard deviation to improve model fit and interpretability (see [31]). For quadratic terms we first mean centred and scaled the variable, then squared it and then divided by two (see: [32]). Each model had the number of different individuals a focal weevil copulated with in that day (our proxy for fitness) as the response variable, with date of observation, tree identity, and weevil identity as random effects, and a Poisson error distribution with a log-link. This approach gives fixed effect coefficients that are directly interpretable as selection gradients (see [32]).

To estimate direct and social selection, in our first model we fitted individual body size and the mean body size of all other individuals of the same sex on the same tree in that day as predictors. The latter term specifically excludes the focal individual from the calculation of the mean [7,14,33]. We also included quadratic versions of these terms to determine whether social selection was non-linear. We included sex as a fixed effect, and the interactions between sex and both focal and rival body size for both linear and quadratic terms to test whether males and females experienced different selection. Females were set as the default sex and so the interaction was modelled as the difference between males and females. We evaluated the “clarity” (see [34]) of the effect of all fixed effects using Wald χ^2^ tests with type II sum of squares using the *Anova* function within the R package “cars” [35]. The degrees of freedom were 1 for all tests unless a subscript is given stating otherwise.

To determine under which conditions social selection is strongest we then fitted a series of models. We used the same starting model as above except we did not include quadratic terms as they had no clear effect (see Results). For the first model, we included an interaction between focal body size and the mean size of its rivals to determine if smaller individuals experienced weaker social selection than larger individuals. We also included the three-way interaction between sex, focal body size, and rival body size, to see if males and females differed in this relationship. As males of only smaller sizes (typically under 40mm, see [27]) may engage in “sneaky” copulations, we also fitted a model where sex was a three-level categorical variable, either “female”, “male over 40mm”, or “male 40mm or under”, and retained the interactions between this new variable and both focal and rival body size. We then fitted two models to test which demographic parameters influenced social selection. The first included weevil density (number of weevils observed on the tree on that day) as a fixed effect and its interactions with both focal and rival body size, including the three-way interactions between density, sex, and either focal or rival mean body size. The second model was equivalent to the density model but included sex-ratio (proportion of weevils on the tree on that day that were male) instead of density. In these two models the key terms are the interactions between density/sex-ratio and the mean body size of rivals, as these terms indicate whether the impact of rival body size on focal individual fitness (and so the strength of social selection) increases or decreases with density/sex-ratio (while the interaction between this term and sex indicates whether this effect differs between the sexes or not).

To estimate the overall phenotypic assortment within each sex we calculated the Pearson correlation between the body size of a focal individual and the mean body size of its rivals, where the variables had been mean centred and divided by their standard deviation [14]. Following our detection of density dependent social selection (see Results), we then decided to test whether phenotypic assortment was density dependent. We stress this was a decision made after viewing our initial results and so should be interpreted appropriately. To do this we fitted a linear model with the mean body size of same-sex rivals as the response variable, the body size of a focal individual, the density of weevils on the tree, the focal individual’s sex, and all two- and three-way interactions between these variables as fixed effects. We also included date, weevil identity, and tree identity as random effects. The response variable and all continuous predictor variables were mean centred and divided by their own standard deviation. The key term here is the interaction between density and the body size of the focal individual, as this indicates whether the relationship between the focal individual and the mean body size of its rivals changes with density (while the interaction between this term and sex indicates whether this effect differs between the sexes or not).

## Results

There was no linear or quadratic social selection in either sex when not taking into account variation in body size, density, or sex ratio (linear social selection = 0.479, se = 0.563, χ^2^ = 0.029, p = 0.864; sex interaction = −0.494, se = 0.591, χ^2^ = 0.700, p = 0.403; quadratic social selection = 0.485, se = 0.498, χ^2^ = 0.268, p = 0.604; interaction = −0.674, se = 0.527, χ^2^ = 1.633, p = 0.201). As previously found [25] both sexes were under approximately equal positive linear direct selection for body size (linear direct selection = 0.232, se = 0.382, χ^2^ = 11.477, p < 0.001; sex interaction = 0.038, se = 0.391, χ^2^ = 0.009, p = 0.923; quadratic direct selection = −0.470 se = 0.669, χ^2^ = 1.551, p = 0.213, sex interaction = 0.395, se = 0.671, χ^2^ = 0.347, p = 0.556). The strength of social selection did not depend on the size of the focal individual for either sex (focal and rival body size interaction = −0.200, se = 0.368, χ^2^ = 1.781, p = 0.182; sex interaction = 0.093, se = 0.380, χ^2^ = 0.060, p = 0.806), nor was it different among different classes of male (contrast between female and large male = −0.197, se = 0.218, contrast between female and small male = −0.014, se = 0.204, χ^2^_2_ = 1.262, p = 0.532). Direct selection also did not differ among different classes of male (contrast between female and large male= −0.311, se = 0.180, contrast between female and small male= −0.390, se = 0.207, χ^2^_2_ = 4.087, p = 0.532). Social selection was density dependent; at higher densities it was clear and negative for both sexes (rival body size and density interaction = −0.563, se = 0.301, χ^2^ = 4.340, p = 0.037, sex interaction = 0.358, se= 0.370, χ^2^ = 0.934, p = 0.334), indicating that larger rivals reduce an individual’s fitness, but only at high densities (Figure 1). Direct selection was not dependent on density for either sex (focal body size and density interaction = −0.043, se = 0.146, χ^2^ = 2.170, p = 0.141; sex interaction = 0.125, se = 0.154, χ^2^ = 0.652, p = 0.419). Sex-ratio did not influence social selection (rival body size and sex-ratio interaction = −0.158, se = 0.156, χ^2^ = 0.060, p = 0.806; sex interaction = 0.175, se = 0.174, χ^2^ = 1.006, p = 0.316) or direct selection in either sex (focal body size and sex-ratio interaction = −0.006 se = 0.167, χ^2^ = 1.209, p = 0.272; sex interaction = 0.070, se = 0.176, χ^2^ = 0.159, p = 0.690;). Full results from each model are reported in the supplementary materials (Tables S1-5).

**Figure 1.**
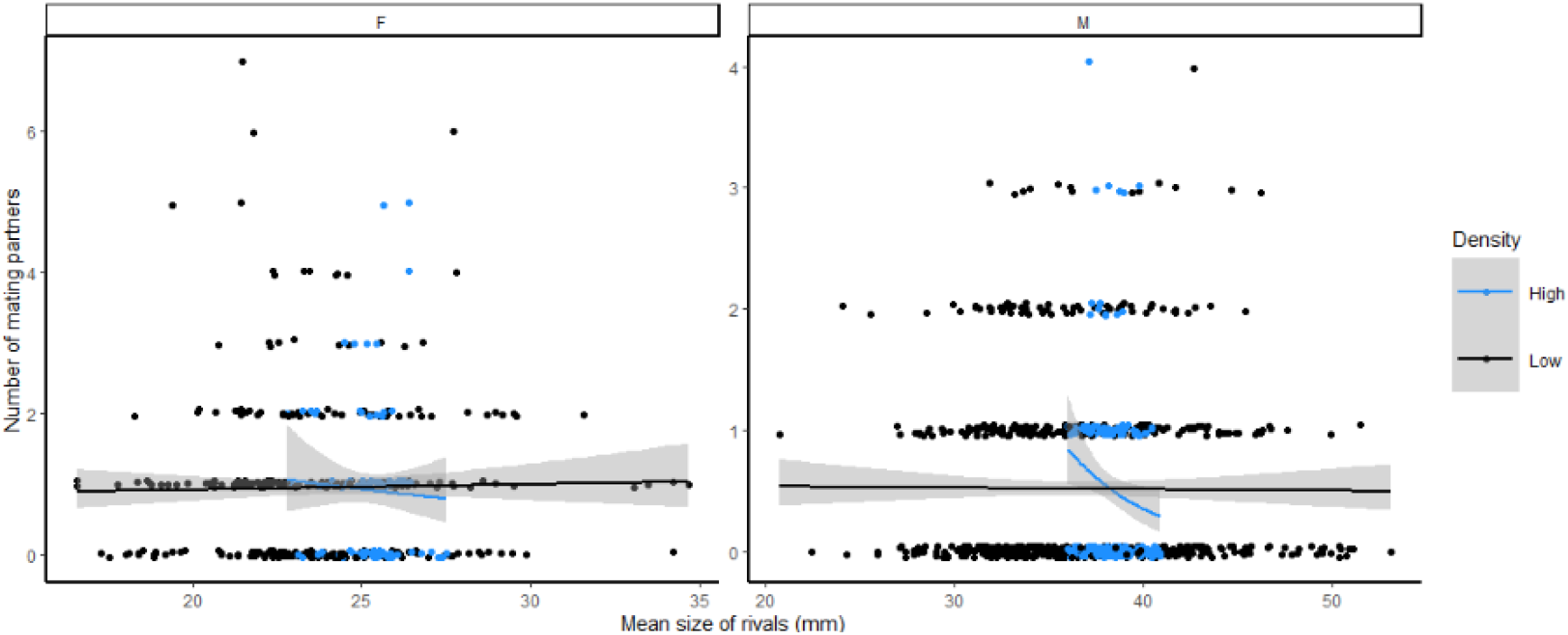
The strength of social selection was density dependent, becoming more negative at high densities (more than 40 weevils per tree, light blue) compared to low densities (40 or fewer weevils per tree, black). This was true for both females (left panel) and males (right panel). Note that we analysed density as a continuous variable, but we have used a categorical representation when plotting for ease of viewing.

Phenotypic assortment overall was near zero for both females (r_females_ = 0.066, t = 1.336, df = 406, p = 0.182) and males (r_males_ = 0.053, t = 1.521, df = 824, p = 0.129). Our subsequent test of whether phenotypic assortment was density dependent revealed that, for both sexes, assortment switched from being positive to negative as densities increased (focal body size = 0.012, se = 0.064, χ^2^ = 18.800, p < 0.001; focal body size and density interaction = −0.021, se = 0.057, χ^2^ = 4.564, p = 0.033; sex interaction = 0.056, se = 0.059, χ^2^ = 0.915, p = 0.339; Figure 2). However, at no density was it especially far from zero, and therefore social selection does not ever greatly alter the total selection differential (Figure 3).

**Figure 2.**
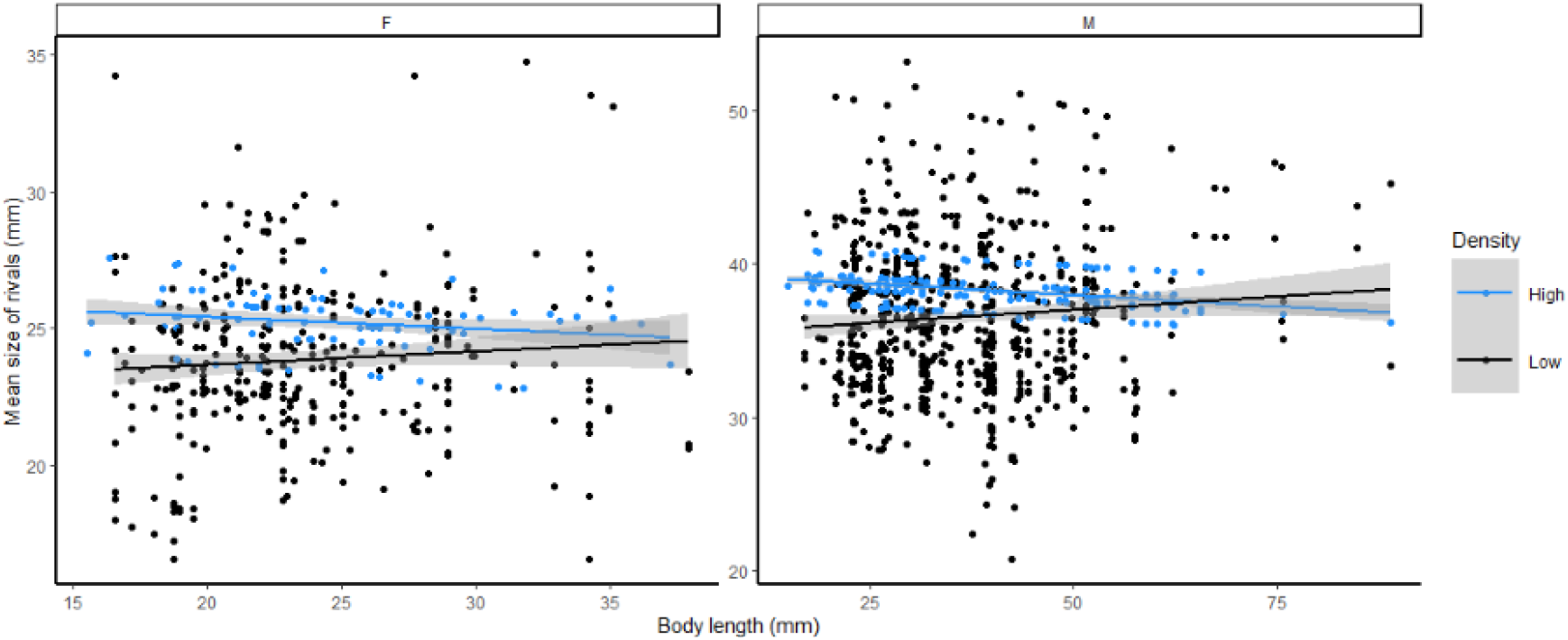
Phenotypic assortment was density dependent, becoming more negative at high densities (more than 40 weevils per tree, light blue) compared to low densities (40 or fewer weevils per tree, black). This was true for both females (left panel) and males (right panel). Note that we analysed density as a continuous variable, but we have used a categorical representation when plotting for ease of viewing.

**Figure 3.**
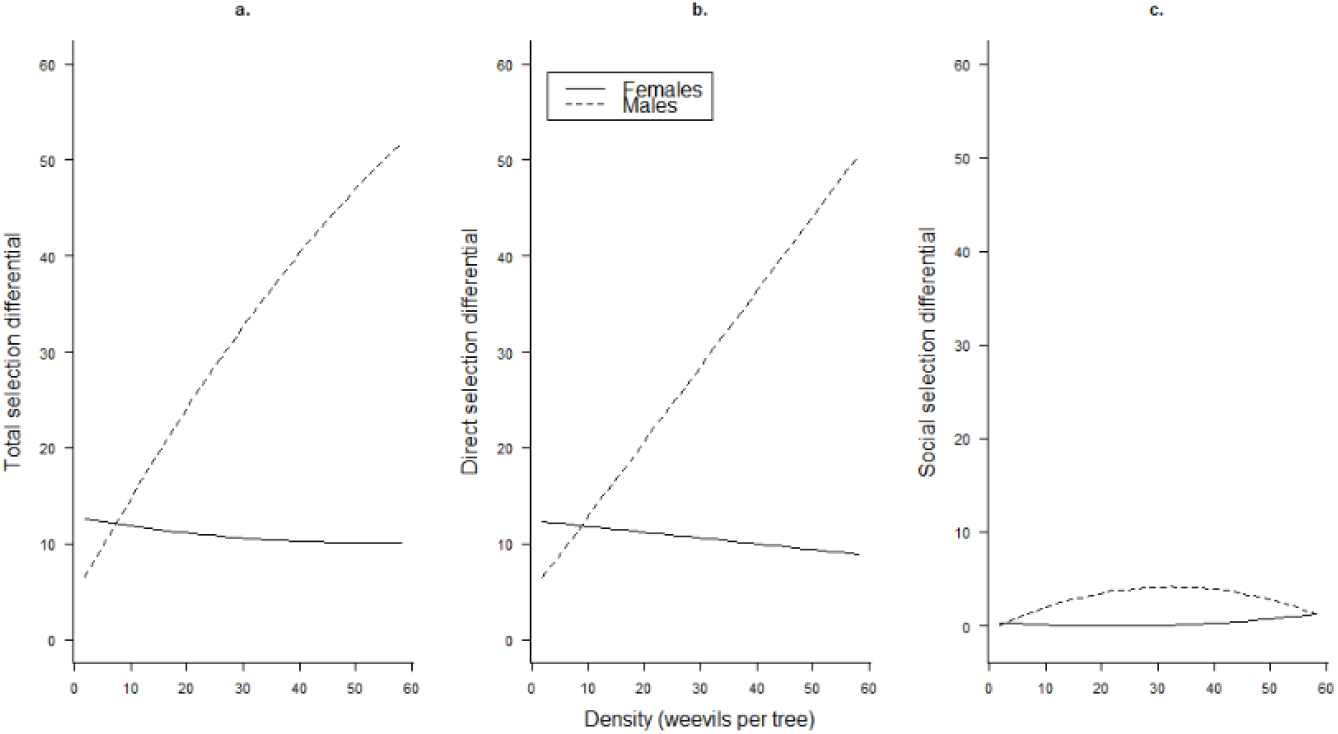
Plots showing how the total selection differential (a.), and its components the direct selection differential (b.) and the social selection differential multiplied by phenotypic assortment (c.) change with density for females (solid lines) and males (dashed lines). Note that direct selection was not clearly density dependent for either sex, but the high phenotypic variance in male body size means small changes in the selection gradient have large consequences for the selection differential. While social selection became more negative with increased density, phenotypic assortment changed from positive at low densities to negative at high densities and was always near zero, meaning the contribution of social selection to the total selection differential was always small.

## Discussion

We estimated the strength of social selection across a range of contexts for both male and female giraffe weevils. In contrast to our predictions, we found that social selection was typically absent, although it was clearly negative at high densities, a result in line with our predictions. An increase in the strength of negative social selection as densities increase is consistent with the idea that weevils are engaging in higher competition for access to mates. Interestingly, this is true for males and females, and so females experience reduced mating success at high densities when there are large females on the same tree. The mechanism for this social selection in females remains to be explored, although there is some evidence larger males prefer to mate with larger females [25], which might lead to fewer matings for smaller females. Another possibility is that at high densities males are spending more of their time fighting other males, leaving less time to copulate with females, resulting in choosier males to the detriment of small females sharing trees with large females. However, as phenotypic assortment was always close to zero the social selection we observed will only make very small contributions to overall selection. At low densities, social selection is weakly positive or absent and so will only very slightly increase total selection (which is positive) due to the positive assortment, while at high densities social selection will also slightly increase overall selection as both it and assortment are negative. Despite this effect, ultimately it seems that direct selection, in interaction with costs and benefits stemming from natural selection [25], will govern the evolution of body and weapon size in giraffe weevils

Our results are consistent with several previous studies on direct and social selection for body size and related traits (see also [7] for a list of direct and social selection in other types of traits). Formica et al. [26] found positive direct selection and negative social selection for body size (when using mating success as a proxy for fitness, but this is not true when using survival) in fungus beetles (*Bolitotherus cornutus*), matching our result for high densities. Similarly, Tsuji [36] and Santostefano et al. [15] found positive direct selection and negative social selection for body size in an ant (*Pristomyrmex pungens*) and in male chipmunks (*Tamias striatus* ; but only in summer, and never for females), respectively. These results have also been repeated in plants, in both Arabidopsis thaliana [37] and sea rocket (*Cakile edentula*) [38] positive direct selection and negative social selection for size has been detected, although in sea rocket this is only true at low densities, while both selection gradients are reversed at medium and high densities. Other studies however either find positive direct selection and either positive or mixed social selection for size or growth rate (black-throated blue warblers, *Setophaga caerulescens*, [39]; North American red squirrels, *Tamiasciurus hudsonicus*, [16]; great tits, *P. major*, [40]; *Silene tatarica* [41]), while Stevens *et al.* [42] found both direct and social selection for size to be negative in Jewelweed (*Impatiens capensis*). Therefore, while opposing direct and social selection for body size may be more common than any other situation, consistent with size-based competition for limited resources which are key for fitness (such as food or members of the time-limited sex), it is by no means the rule. More estimates of direct and social selection need to be accumulated before we can start to identify general rules.

Sex-ratio had no effect on social or direct selection. No effect on social selection surprised us given we assumed social selection represents competition for mates, which should be stronger in males when an aggregation is more male-biased, and potentially vice versa for females. Sex ratio varied from 0-1.0 (median = 0.66, 25% quantile = 0.61, 75% quantile = 0.71) so we do not think it is a lack of variation in our dataset preventing us from finding a pattern. Possibly, many males on a tree on any given day are not participating in the competition for mates, therefore rendering the measure of sex-ratio uninformative. We also found a focal individual’s body size (either measured continuously or where males were split into “small” and “large” male either side of 40mm) did not influence the impact of rivals on fitness. We had expected smaller males to be less severely affected by large rivals, as they are able to obtain matings by switching from a female-defence strategy to “sneaking” copulations with females guarded by large males [27]. However, given we only detected any negative effect of larger rivals at high densities, we might have to focus on high densities to look for our predicted pattern, and the current dataset does not contain enough samples of trees with a high density of giraffe weevils to do this. While correlational direct selection has received some attention [43], we possess very limited information about which traits of individuals influence the strength of social selection (but see [16] for an interaction between sex and multilevel selection on birth date in North American red squirrels, *T. hudsonicus*). Beyond body size, certain behavioural traits, such as sensitivity or susceptibility, might modulate how strongly an individual is influenced by rivals, but this remains to be tested.

We found phenotypic assortment was typically near zero, although was more positive at low densities and more negative at high densities, for both sexes. Due to this near-zero assortment, social selection will only ever contribute a small amount to total selection. Limited phenotypic assortment is consistent with individuals mostly randomly aggregating on trees without respect to the body size of other individuals on the tree. In giraffe weevils, assortment by body size has been observed in mating pairs [25], but this pattern could emerge following arrival at trees rather than before. We did see negative phenotypic assortment at high densities, which is consistent with large individuals avoiding large rivals at high densities, when the fitness consequences of interacting with them is strongest. However, this effect is relatively weak, probably due to individuals of all sizes benefiting from avoiding large rivals, in which case no strong assortment can arise.

Although estimates of phenotypic assortment have been accumulated in the literature and are often positive, they tend not to be specifically measures of the interactant covariance (the covariance between an individual’s traits and the mean trait value of those it interacts with), the key parameter for models of social selection [14]. For example, Farine and Sheldon [44] estimated positive assortment on lay date in great tits (*P. major*) but used a social network measure of assortment which underestimates the true interactant covariance substantially [14]. If positive phenotypic assortment is indeed common, then social selection will often contribute to total selection, and if social selection is typically in the opposite direction to direct selection [7], will therefore tend to reduce overall selection. Formica *et al.* [26] estimated the interactant covariance for body size in aggregations of forked fungus beetles (*B. cornutus*) and found a negative covariance. This would then end up causing negative social selection for body size to increase the magnitude of the overall positive selection for body size. In contrast, while Santostefano *et al.* [15] found a negative covariance among female chipmunks (*T. striatus*) for body mass, they found no covariance among males for body mass. Since social selection was only present in males, social selection would not contribute to overall selection in either sex. In summary then, while we may expect social selection to weaken overall selection, evidence from systems where both social selection and phenotypic assortment have been estimated suggests that it often does not contribute at all. Further, a lack of estimates of how phenotypic assortment changes with key demographic parameters such as density prevents us from understanding whether there are some contexts social selection does contribute to total selection. Given both direct and social selection can also vary with conditions, context-dependent phenotypic assortment raises the possibility that evolution can have very different outcomes in different environments, but we lack the data to assess this suggestion.

Overall, we have contributed to our knowledge of how selection operates in wild animals. As predicted, social selection was in the opposite direction to direct selection and was stronger at high densities. However, social selection was not clearly different from zero in average conditions and did not vary with sex-ratio or the size of the focal individual. Further, although phenotypic assortment changed with density it was rarely far from zero, indicating that social selection will have a limited contribution to overall selection even at high densities. Therefore, despite its predicted importance, social selection will only have a minor impact on the evolutionary change of body size in New Zealand giraffe weevils.

## Data accessibility

The data used here have previously been made publicly available, see: [28]. We have chosen to provide copies of the exact spreadsheets and the R code used to create the dataset, analyse the data, and produce all figures, as supplementary materials for ease of access for reviewers. Upon acceptance we will make these files available in Dryad or another suitable public repository.

## Acknowledgements

We thank John Staniland and Forest and Bird Waitakere for continuously supporting our research at Matuku Reserve. Data collection was made possible by many volunteers, especially Jessica Le Grice, Robin Le Grice and Stephen Wallace. We have no competing interests.

## Authors’ contributions

DNF developed the main ideas for the manuscript, analysed the data, and lead the writing of the manuscript. RLG collected and curated the data. CJP helped collect the data and design the study, provided assistance during the statistical analysis, and contributed to the writing. All authors gave final approval for publication and agree to be held accountable for the work performed therein.

